# Establishing label-free quantitative X-ray dose-response profiling by Holo-Tomographic Flow Cytometry

**DOI:** 10.64898/2026.01.12.699031

**Authors:** Daniele Pirone, Chiara de Vita, Rocco Mottareale, Giusy Giugliano, Gennaro Giordano, Simonetta Grilli, Vittorio Bianco, Lisa Miccio, Pasquale Memmolo, Marco Durante, Mariagabriella Pugliese, Pietro Ferraro

## Abstract

Flow cytometry (FC) offers multiparametric analysis capabilities that can quantify cellular damage after exposure to cytotoxic agents. Here, we present a comprehensive study establishing a label-free quantitative X-ray dose-response profiling using a novel FC modality based on 3D Quantitative Phase Imaging, termed Holo-Tomographic Flow Cytometry (HTFC). This approach enables fully label-free 3D refractive index (RI) measurements, allowing detailed and quantitative characterization of the biophysical properties and morphology of living cells exposed to X-rays. By analyzing datasets of 3D RI tomograms from cells irradiated at graded doses, we identify intracellular biophysical markers that define a robust X-ray dose-response curve. Validation against standard clonogenic survival assays on three model cancer cell lines reveal a high correlation (>90%). HTFC not only eliminates labeling and operator bias but also markedly reduces experimental time from 1-2 weeks to 24 hours, offering a fully automated and objective readout. While clonogenic survival remains the benchmark for radiosensitivity assessment, our findings establish HTFC as a powerful label-free platform for fast assessment of radiation damage. This technology paves the way for predictive biosensors that can capture patient-specific responses, thereby supporting the transition from conventional, uniform radiotherapy protocols to personalized treatment strategies.

## Introduction

Cell populations are inherently heterogeneous in morphology, size, cell cycle phase, and physiological state [1]. For this reason, the detailed characterization of individual cells has become a major objective in contemporary biomedicine and personalized medicine, which increasingly rely on advanced cytometric platforms for the single-cell analysis [2]. A significant advancement in cytometric analysis has been the integration of Flow Cytometry (FC) [3] with microscopy, leading to Imaging Flow Cytometry (IFC). Over the past decades, IFC has enabled substantial progress in cell biology studies [4], rapidly establishing itself as a reference method in diverse biomedical fields, including cancer research [5]. IFC simultaneously records bright-field, dark-field, and fluorescence images from individual cells as they move in a microfluidic stream, thus providing a high-throughput acquisition of spatially resolved data. The resulting large datasets allow extraction of multiple morphological and functional parameters at the single-cell level. Among its emerging applications, IFC has recently shown great promise in biodosimetry [6-8], a field devoted to quantifying biological responses to ionizing radiation for the estimation of absorbed doses and evaluation of radiation-induced cellular damage [9]. For example, IFC has been applied to evaluate radiation response in lymphocytes from umbilical cord blood and cancer patients [10] or to enable high-throughput quantification of DNA double-strand break repair kinetics through γ-H2AX analysis [11]. Despite its significant advantages, IFC still faces several drawbacks that limit its broader adoption in biomedical research. The technique remains inherently fluorescence-dependent, operator-sensitive, and often time-consuming and costly. It requires prior knowledge of appropriate exogenous markers, which can affect measurement accuracy due to photobleaching and may introduce phototoxic effects that alter cellular physiology. Once labelled, cells cannot be reused for downstream analyses. Most critically, IFC provides limited access to the intrinsic biophysical properties of cells, as it primarily captures morphological and fluorescence intensity features rather than quantitative physical information.

Therefore, label-free approaches have recently attracted growing interest, as they allow the study of cellular processes without relying on exogenous dyes. For instance, direct mass spectrometry has been applied to investigate bystander effects in chondrocytes co-cultured with chondrosarcoma cells irradiated with X-rays or carbon ions [12], Raman micro-spectroscopy has been used to assess the impact of X-ray irradiation on nuclear and membrane regions of single neuroblastoma cells [13], and, more recently, lab-on-a-chip impedance spectroscopy has been employed to monitor radiotherapy responses even in 3D spheroids [14]. Among label-free microscopy methods [15], Quantitative Phase Imaging (QPI) has emerged as a particularly powerful tool over the past decades [16]. Typically, QPI exploits the interferometric properties of Digital Holography (DH) to reconstruct Quantitative Phase Maps (QPMs) of individual cells [17,18]. In a QPM, both cell morphology and the spatial distribution of the refractive index (RI) are encoded in a 2D image. Thus, unlike fluorescence images, QPMs provide more distinctive information on the cellular state [19], since the RI is a fundamental biophysical parameter [20,21]. DH has been proposed as a label-free approach for the non-invasive, quantitative assessment of the morphological changes of γ-irradiated human mesenchymal stem cells and periosteal cells [22], X-irradiated breast cancer cells [23], and X-irradiated urothelial bladder carcinoma cells [24]. Moreover, combined to FC, we have recently demonstrated that DH can measure the effects of X-ray radiations on single flowing lymphocytes and classify the irradiation dose by machine learning tools [25]. The most recent advancement in label-free bioimaging is the 3D extension of QPI, known as Holographic Tomography (HT), which opens unprecedented opportunities by providing access to the full RI volumetric distribution of single cells [26]. Among the various HT configurations [27,28], we have developed Holo-Tomographic Flow Cytometry (HTFC), which integrates the label-free, quantitative, and 3D capabilities of HT with the single-cell analysis typical of IFC [29-35].

Here, by exploiting the unique capabilities of HTFC, we present for the first time a label-free quantitative X-ray dose-response profiling obtained using a 3D QPI technique. Thus, we correlate 3D biophysical markers with cancer cell responses to defined X-ray irradiation strategies and their effectiveness. In the FC configuration, we acquire comprehensive datasets of 3D RI tomograms from single cells exposed to different X-ray irradiation doses. An *ad hoc* statistical segmentation algorithm [29] is then applied to extract nucleus-specific information from stain-free, quasi-spherical cells, thereby faithfully reproducing the intracellular organization under suspension conditions. For each dose, we derive quantitative single-cell biomarkers, which enable us to evaluate the effects of X-ray irradiation across the entire cell population and, ultimately, to construct a label-free dose-response curve. By analysing clonogenic assays in three different model cancer cell lines, we discover an intriguing duality between this label-free curve and gold-standard dose-response profiles, as we observe a strong correlation (>90%) between the two approaches.

Beyond its label-free nature, the proposed method provides other significant advantages. Firstly, the time required to obtain a dose-response curve is drastically reduced, as the HTFC experiment can be completed 24 hours after irradiation, whereas clonogenic assay readouts require 1-2 weeks. Second, clonogenic assay results are based on a human colony counting, thus they are prone to a bias introduced by the operator; instead, HTFC measurements are inherently objective, as they provide quantitative physical information about each cell. Moreover, clonogenic assays provide an indirect measure of radiosensitivity by quantifying the fraction of cells that retain the ability to proliferate after irradiation, thus reflecting only the ultimate cell death outcome. In contrast, HTFC allows us to directly evaluate the radiation-induced alterations in viable cells before they reach the death state, offering in principle a dynamic view of those biophysical and morphological changes that typically precede the loss of clonogenic potential. We demonstrated in a previous work that label-free 2D QPI measurements can classify irradiation doses and capture their effects on living cells [25]. Starting from that preliminary result, here we exploit a 3D QPI technique such as HTFC to reveal distinctive signatures of radiosensitivity and to establish a label-free biosensor [36] capable of predicting tumour responses to ionizing radiation in a rapid, automatic, and objective manner. This approach could open new routes on personalization of radiotherapy (RT) treatments, which remain a cornerstone of multidisciplinary cancer care [37]. In fact, current RT protocols still rely on a one-size-fits-all principle, yet it is well recognized the importance for tailoring treatment to individual patient characteristics and selecting the most appropriate RT protocol as patient responses to radiation vary widely [38].

## Results

### Extraction of 3D biophysical markers to characterize the HTFC dataset at different X-ray irradiation doses

Using the HTFC experimental system, the neuroblastoma SKNBE2 cell line was analysed 24 hours after X-ray irradiation at different doses. The primary objective was to investigate the cellular response to a heterogeneous spectrum of X-ray exposures under the following conditions: low doses (0.25 and 0.5 Gy), intermediate doses (1 and 1.5 Gy), and the standard fractionated dose typically used in clinical radiotherapy (2 Gy). For each cell, the corresponding 3D RI tomogram was reconstructed, yielding the dataset summarized in Table S1 corresponding to 590 SKNBE2 cells. Figure 1 shows the typical HTFC reconstructions obtained for different SKNBE2 cells 24h after the exposure to 6 different X-ray irradiation doses, from 0 Gy up to 2 Gy. In particular, Figure 1(a) shows the representative central RI slices of the irradiated cells.

**Figure 1.**
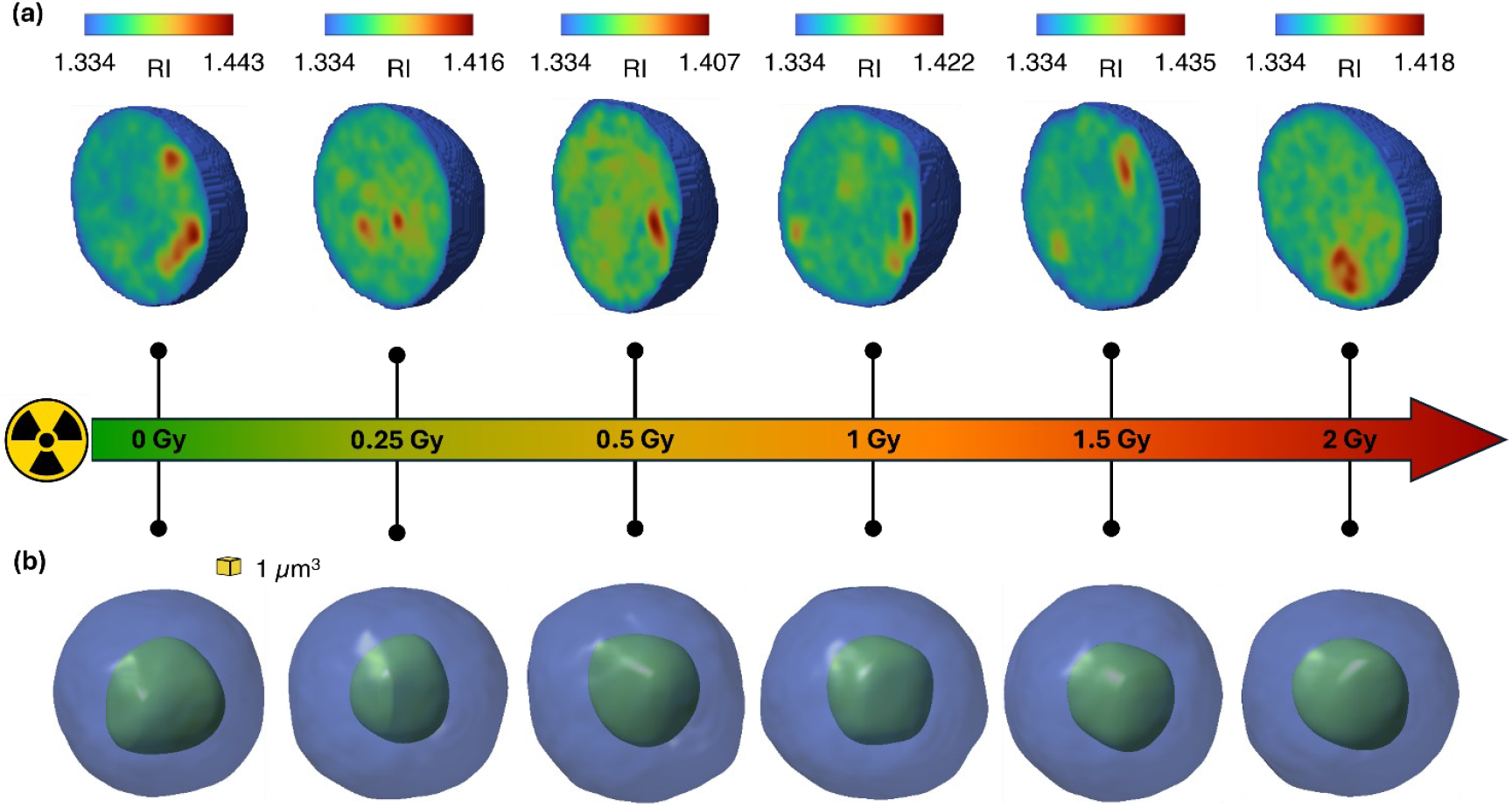
HTFC reconstructions of different neuroblastoma SKNBE2 cells 24 h after the exposure to different X-ray irradiation doses. **(a)** Central slice of the 3D RI tomograms. **(b)** Isolevels representation of the 3D RI distribution in (a), with the CSSI-segmented nucleus (green) within the cytoplasm (blue).

In label-free techniques such as QPI, the main limitation is the lack of intracellular specificity, since no marker is available to distinguish organelles from their background. Typically, intracellular RI contrast alone is insufficient to clearly differentiate subcellular structures [39]. To address this issue, advanced computational strategies have recently been developed to enhance intracellular specificity [40], including virtual staining methods based on deep learning [41]. In the case of HTFC, this challenge is even greater due to the suspended condition of single cells. To overcome it, we have previously demonstrated a Computational Segmentation based on Statistical Inference (CSSI) approach [29,30], which identifies organelles by exploiting the statistical similarity among voxels within their 3D RI volume. Since it is well established that ionizing radiation affects nuclear morphology [42], we applied the CSSI algorithm to segment the nucleus in the 3D RI tomograms of each SKNBE2 cell. Examples of segmented nuclei are reported in Fig. 1(b).

Building on the retrieved nuclear information, we carried out a multiplexed quantitative characterization at the single-cell level by extracting 39 biophysical markers related to the whole cell, nucleus, and cytoplasm (summarized in Table S7). For each compartment, we calculated seven features: RI mean, RI standard deviation, RI coefficient of variation, RI maximum, RI interquartile range, volume, and dry mass. The RI coefficient of variation was defined as the ratio between the RI standard deviation and the RI mean, while the dry mass represents the mass of the object excluding its aqueous content [16]. For each of these seven features, we also calculated the ratio between nuclear and cytoplasmic values. Moreover, for both the whole cell and the nucleus, we evaluated three morphological descriptors: sphericity, solidity, and extent. Sphericity was defined as the ratio of the surface area of a sphere with the same volume to the object’s surface area, solidity as the ratio between the object’s volume and the volume of its convex hull, and extent as the ratio between the object’s volume and the volume of its bounding box. For each of these three descriptors, we additionally computed the ratio between nuclear and whole-cell values. Finally, we introduced two nucleus-to-cell features: the nucleus–cell normalized distance, defined as the Euclidean distance between the centroids of the nucleus and the cell normalized to the equivalent cell radius (i.e., the radius of a sphere with the same volume as the cell), and the nucleus–cell normalized solid angle, defined as the solid angle subtended by the nucleus from the cell centroid, normalized to 4π.

We ranked the 39 biophysical markers according to their ability to discriminate among the different irradiation doses using the ReliefF algorithm with 10 nearest neighbours [43]. As shown in the ranking reported in Fig. 2(a), the RI statistics of the various intracellular compartments (cell, nucleus, and cytoplasm) emerged as the most informative features, followed by their morphological features (i.e., sphericity, solidity, extent, and volume). This highlights the robustness of the selected feature set, which combines morphological descriptors (size and shape) with RI-based biomarkers calculated at both the whole-cell and intracellular levels, providing a meaningful characterization of the HTFC dataset across irradiation conditions. In Fig. 2(b), boxplots and corresponding one-sided violin plots are presented for some of the highest-ranked features. However, all the selected features display an oscillatory behaviour across the irradiation doses compared to the control condition, indicating that none of them individually shows a monotonic trend suitable to serve as a reliable dose–response indicator. For instance, the cell RI maximum (1st) decreases from 0 Gy to 1.5 Gy and then increases again at 2 Gy; the nucleus RI standard deviation (5th), cell sphericity (6th), and cytoplasm RI interquartile range (14th) increase at 0.25 Gy, decrease at 0.5 Gy and 1 Gy, and rise again at 1.5 Gy and 2 Gy; the cell volume (16th) and nucleus dry mass (24th) exhibit alternating oscillations across the dose range.

**Figure 2.**
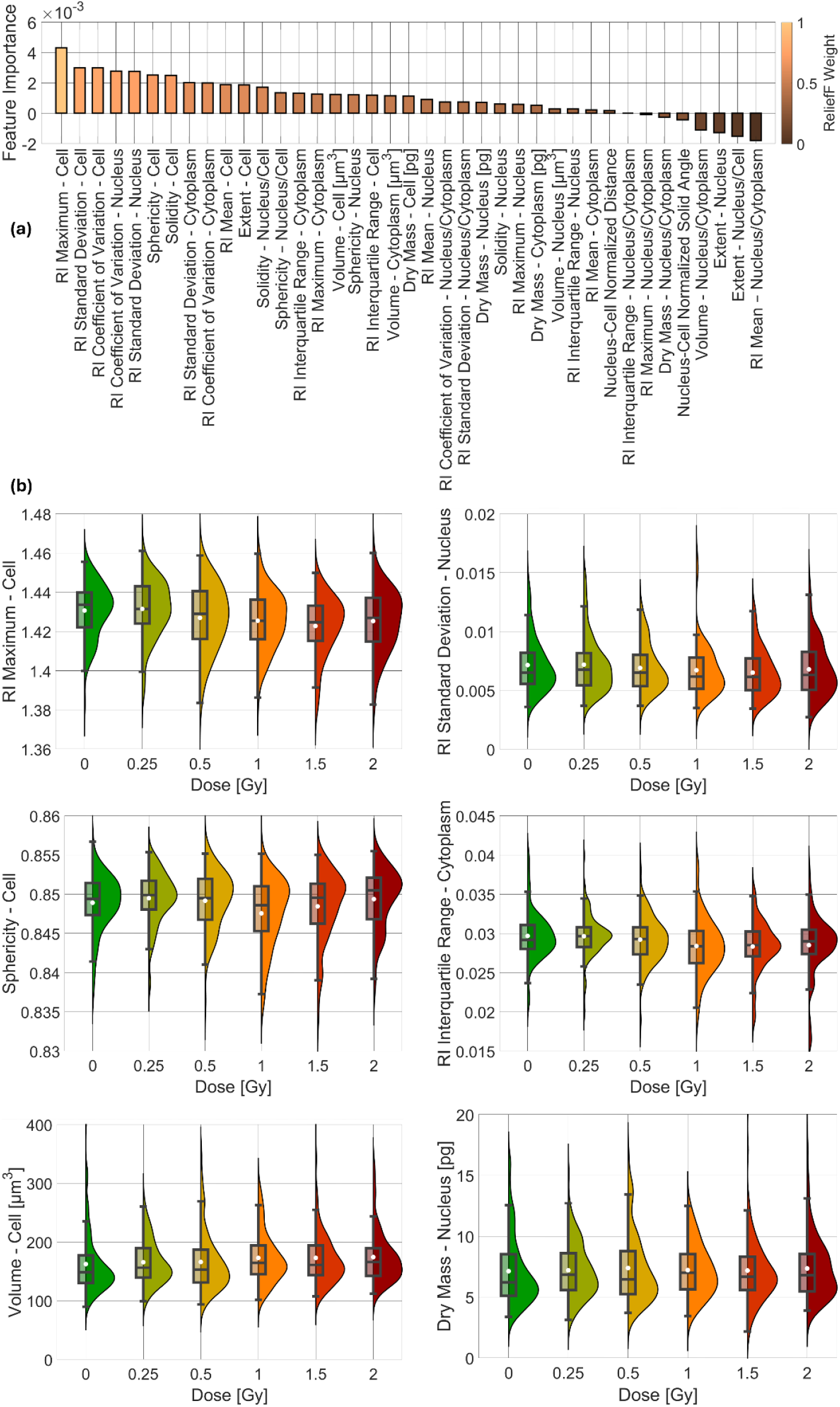
Biophysical markers used to characterize the HTFC dataset of neuroblastoma SKNBE2 cells 24 h after exposure to different X-ray irradiation doses. **(a)** Feature ranking based on the ReliefF weights. **(b)** Boxplots and corresponding one-sided violin plots related to some of the highest-ranked features in (a). In each box, the white dot represents the mean value, the line inside represents the median, the lower and upper box limits correspond to the first and third quartiles, and the whiskers extend to the furthest values within the non-outlier range.

### Definition of a label-free quantitative X-ray dose-response curve

We analysed the HTFC multiparametric redout by means of a principal component analysis (PCA) using all 39 biophysical markers as input [44]. This allowed each cell to be represented in the space defined by the first two PCA components, as displayed in Fig. 3(a). As our goal was to define a dose-response curve, we performed pairwise comparisons of each irradiated cell population against the control population in the 2D PCA space, as shown in Figs. 3(b-f). Notably, the cluster of cells irradiated at a given X-ray dose shifts progressively further from the control cluster as the dose increases. However, to quantify this shift, it is necessary to account not only for the Euclidean distance between cluster centroids but also for the dispersion of each cluster. Accordingly, Fig. 3 also displays the covariance ellipse for each cluster, whose shape and orientation are determined by the eigenvectors and eigenvalues of the cluster covariance matrix. These ellipses provide a visual representation of the data’s dispersion and correlation along the principal axes. For multivariate normal distributions, each ellipse corresponds to the 1-standard deviation contour around the cluster centroid. For example, the 0.25 Gy population in Fig. 3(b) exhibits the narrowest dispersion, while the 1 Gy population in Fig. 3(d) shows the largest one. In contrast, the 0.5 Gy, 1.5 Gy, and 2 Gy populations in Figs. 3(c,e,f), respectively, display a dispersion comparable to that of the control group.

**Figure 3.**
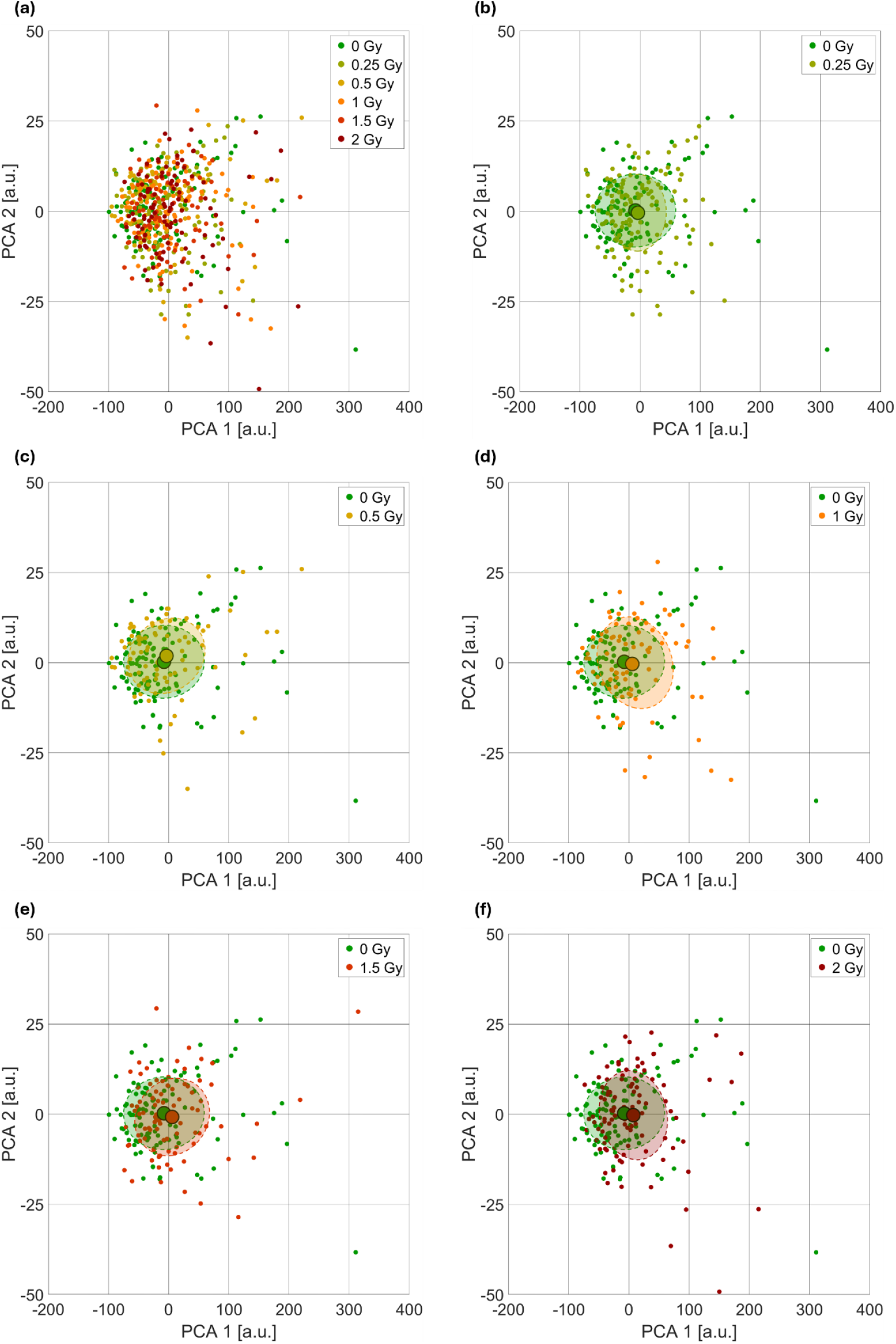
Representation in the 2D PCA space of the neuroblastoma SKNBE2 cells exposed to different X-ray doses according to their HTFC-based biophysical characterization. **(a)** Comparison between the control population and the irradiated populations. **(b)** Comparison between the control population and the population irradiated at 0.25 Gy. **(c)** Comparison between the control population and the population irradiated at 0.5 Gy. **(d)** Comparison between the control population and the population irradiated at 1 Gy. **(e)** Comparison between the control population and the population irradiated at 1.5 Gy. **(f)** Comparison between the control population and the population irradiated at 2 Gy. For each cluster, the centroid and the covariance ellipse are highlighted.

Therefore, to quantify differences among cell populations, we computed the distance of each irradiated cluster from the control cluster in the 2D PCA space using the Fisher Discriminant Ratio (FDR) [45]. In fact, the FDR takes into account not only the separation between cluster centroids but also the dispersion within each cluster, providing a robust measure of the overall distinction between populations. Let 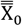 and 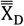 be the PCA matrix of the control population and the population irradiated by a certain dose D, respectively, where the PCA matrix contains the PCA features of all single cells of that population. Thus, 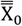 and 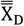 have size N_0_ × d and N_D_ × d, respectively, with d being the dimensionality of the PCA (i.e., 2 in this study), and N_0_ and N_D_ being the number of cells in the control population and irradiated population, respectively. We computed the FDR at a certain dose D as

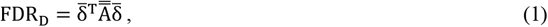

with

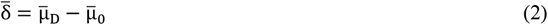

and

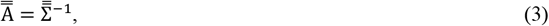

where 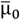 and 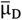 respectively are the mean vectors of 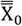 and 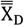 across single cells (size d × 1), and 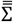 is the total covariance matrix given by the sum of the control covariance matrix 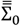 and the irradiated covariance matrix 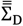.

Moreover, to estimate the uncertainty associated with the FDR_D_ between the multivariate control and irradiated populations, we used analytical error propagation based on a first-order Taylor expansion (also known as Gaussian error propagation) [46]. This method approximates the variance of a function of random variables based on the variances and covariances of its inputs. To propagate the uncertainty in FDR_D_, we consider the uncertainty in the mean vector 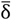 by calculating its covariance matrix as

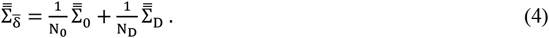

At this stage, we assume to neglect the uncertainty in the covariance matrices 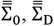, and then 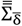. This allows us to treat 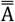 as a constant with respect to the uncertainty propagation. While this introduces a simplification, it enables a tractable analytical expression and remains a reasonable approximation when the number of samples is sufficiently large. Under this assumption, the uncertainty of FDR_D_ is

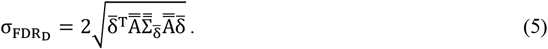

This expression provides a first-order estimate of the uncertainty in the FDR_D_ distance due solely to the variability in the sample means. It is valid under the assumption that the sample means are approximately normally distributed, which is held asymptotically by the Central Limit Theorem. The FDR_D_ distances of the neuroblastoma SKNBE2 population are reported in Table S2 along with their uncertainties 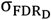.

In summary, at each irradiation dose D, the corresponding FDR_D_ distance quantifies the divergence of the irradiated cell population from the control population. Accordingly, we define the curve formed by the FDR_D_ values vs. the radiation dose D as the divergence score (DS) curve. The DS curve for the neuroblastoma SKNBE2 cell line is shown in Fig. 4(a). We hypothesize that the DS curve can serve as a label-free quantitative X-ray dose-response curve, as it exhibits the typical characteristics expected for radiobiological dose–response relationships. Specifically, the DS increases monotonically with the dose, indicating a progressively larger deviation from the control population as the irradiation intensity rises. The curve originates from a defined baseline (DS = 0) in the absence of irradiation, reflecting the unperturbed state of the control cells. Furthermore, the DS curve exhibits a nonlinear trend with increasing dose, consistent with the saturating and sigmoidal behaviors commonly observed in classical dose-response curves. Taken together, these features suggest that the DS curve not only captures the magnitude of the cellular response to X-ray exposure but also provides a single-cell quantitative readout that mirrors the fundamental properties of conventional dose-response assays. These results strongly support its potential application as a label-free alternative to standard clonogenic assays for evaluating radiation sensitivity.

**Figure 4.**
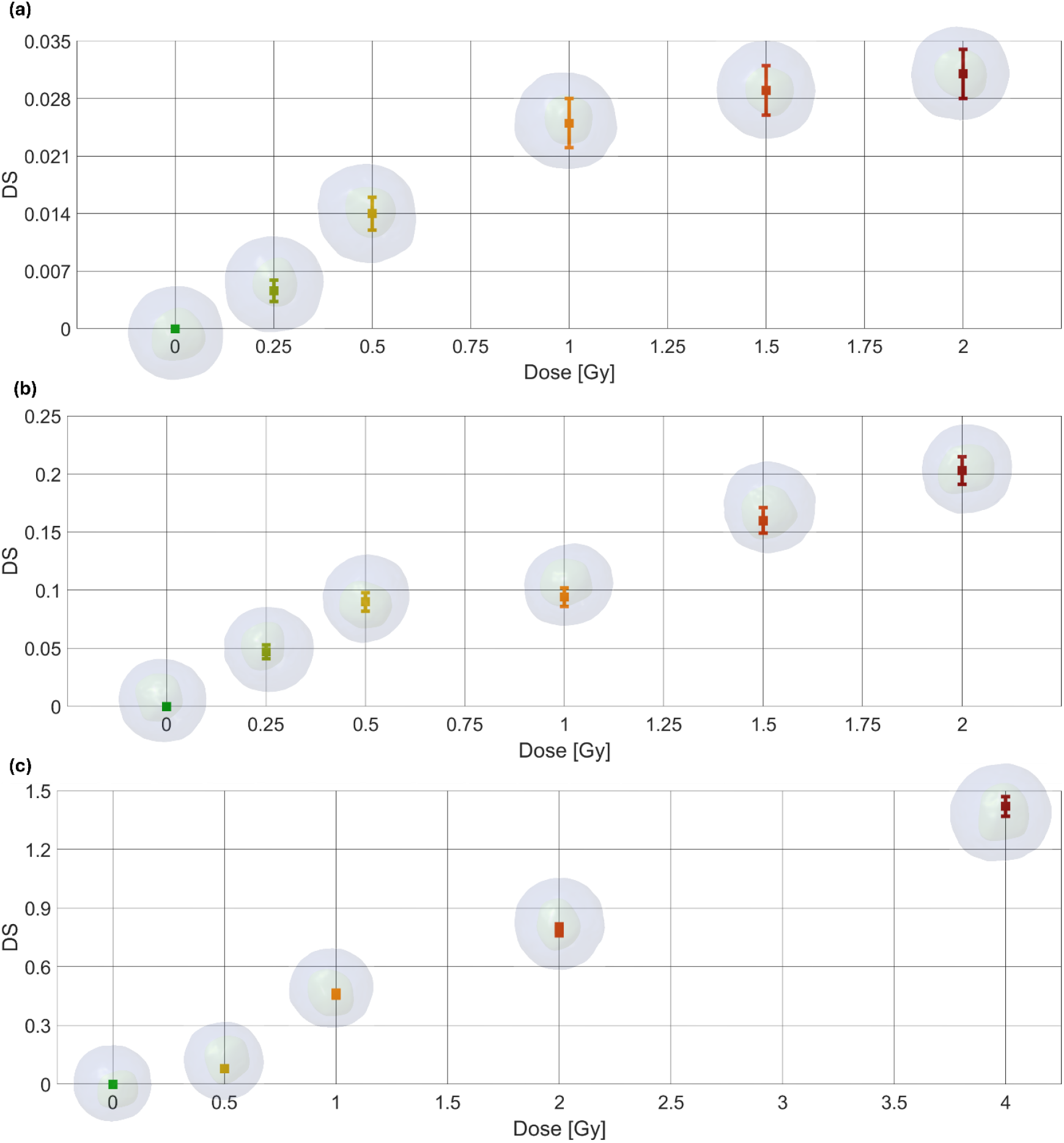
DS curve computed by HTFC to serve as a label-free quantitative X-ray dose-response curve. **(a)** Neuroblastoma SKNBE2 cell line. **(b)** Breast cancer MCF7 cell line. **(c)** Neuroblastoma SHSY5Y cell line.

### Assessment of the Divergence Score as a label-free quantitative X-ray dose-response curve

The clonogenic assay is considered the gold standard in radiobiology [47]. In this assay, single cells are seeded at low density and allowed to grow for 1-2 weeks, during which only cells that survive the radiation insult and preserve their reproductive capacity give rise to colonies. Colonies are then fixed, stained, and manually counted. A parameter commonly employed in radiobiology to assess the response of a cell population to ionizing radiation is the surviving fraction (SF), which is conventionally measured through clonogenic assays. The SF is defined as the ratio between the number of colonies formed by irradiated cells and the number of seeded cells corrected for the plating efficiency. In other words, the SF quantifies the proportion of cells that retain the ability to undergo unlimited proliferation and form macroscopic colonies following radiation exposure. The resulting SF curve, obtained by plotting the SF values against the radiation dose, typically exhibits a decreasing trend confined within the [0,1] interval, reflecting the progressive loss of clonogenic potential with increasing radiation dose.

Hence, here we exploit the SF curve obtained by clonogenic assay (see Table S2) to assess the DS curve as a label-free quantitative X-ray dose-response curve. For the sole purpose of comparing the trend of the DS curve to the trend of the SF, we computed the normalized divergence score (NDS) as

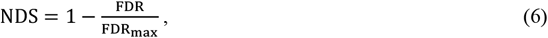

where FDR_max_ is the maximum value of the DS curve in the observed dose range. As reported in Table S2, thanks to the normalization, the NDS value at 0 Gy is forced to be 1 and the NDS value at the highest irradiation dose is forced to be 0, thus reproducing a decreasing trend in the [0,1] interval.

Accordingly, the uncertainty on NDS_D_, denoted as 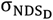, was estimated using standard first-order error propagation. The total propagated uncertainty is given by

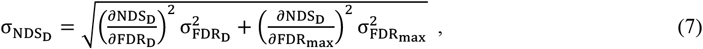

which yields

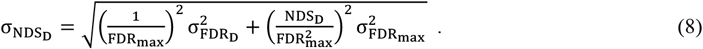

In Figs. 5(a,b), for the neuroblastoma SKNBE2 cell line, we report the SF and NDS curves, respectively. To evaluate the relationship between the two readouts, Fig. 5(c) shows their linear regression analysis by y = m(x − 1) + 1 equation, which yields a slope of 0.73 ± 0.03. This corresponds to a Pearson correlation coefficient 0.99 ± 0.01, confirming a very strong agreement between the trends of the two approaches despite intrinsic differences in their absolute values. In fact, both curves originate from a value of 1 at 0 Gy, reflecting the unperturbed state of the control population. However, while the SF curve progressively decreases with increasing dose and asymptotically approaches 0 (without necessarily reaching it in the dose range considered), the NDS curve is inherently normalized to span the [0,1] interval, with its minimum always equal to 0. This normalization accounts for the divergence-based nature of the metric and explains the systematic offset observed between the two curves.

**Figure 5.**
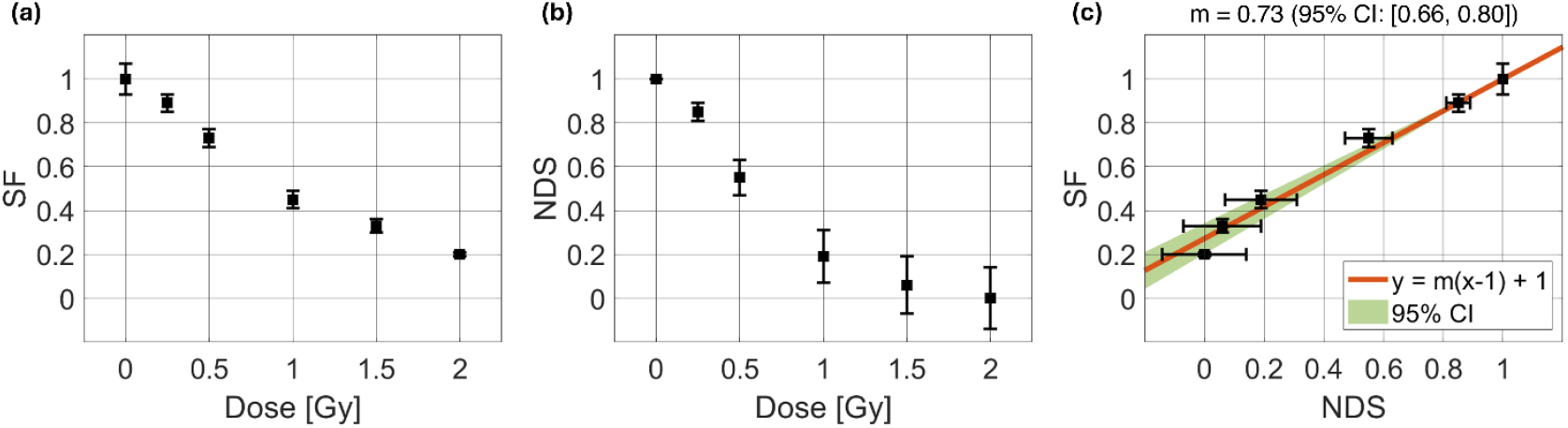
Assessment of the label-free quantitative X-ray dose-response curve for the neuroblastoma SKNBE2 cell line. **(a**,**b)** SF and NDS curves, respectively. **(c)** Correlation between SF curve in (a) and NDS curve in (b).

To further validate our DS dose-response curve, we performed an additional experiment using a different cancer type by analysing breast cancer MCF7 cells. As summarized in Table S3, we collected 528 cells using the same irradiation doses as for the neuroblastoma SKNBE2 experiment. The DS curve for MCF7 cells (see Fig. 4(b)) exhibits the same characteristic features observed for the neuroblastoma SKNBE2 cell line (see Fig. 4(a)), showcasing the reproducibility of the proposed approach across different cell types. The corresponding SF and NDS curves for MCF7 cells are shown in Figs. S1(a,b), respectively (see also Table S4), yielding a slope of 0.65 ± 0.04, corresponding to a Pearson correlation coefficient 0.98 ± 0.03 (see Fig. S1(c)).

Finally, we extended the validation to a different neuroblastoma cell line (i.e., 283 SHSY5Y cells). To test the HTFC system under modified experimental conditions, we adjusted the irradiation interval, which was enlarged and down sampled. In particular, the intermediate doses of 0.25 Gy and 1.5 Gy were omitted, while the higher dose of 4 Gy was included, as summarized in Table S5. The resulting DS curve for the neuroblastoma SHSY5Y cells (see Fig. 4(c)) exhibits a nonlinear monotonic trend, similar to that observed for the neuroblastoma SKNBE2 cells in Fig. 4(a) and breast cancer MCF7 cells in Fig. 4(b), supporting the applicability of the DS curve as a label-free dose-response metric. To further corroborate this observation, we compared the corresponding SF and NDS curves for the neuroblastoma SHSY5Y cell line (see Figs. S2(a,b), respectively; see also Table S6). Regression analysis yielded a slope of 1.12 ± 0.16, corresponding to a Pearson correlation coefficient 0.94 ± 0.11 (see Fig. S2(c)), confirming a strong agreement between the traditional and label-free readouts despite the down sampled irradiation interval and the smaller single cells dataset.

## Discussion

We introduced a novel approach to quantify radiation-induced cellular alterations by establishing a label-free quantitative X-ray dose-response curve from HTFC 3D quantitative single-cell analysis. By exploiting 3D RI tomograms, we extracted a panel of biophysical markers descriptive of both whole-cell and intracellular compartments. Through PCA and FDR, we defined a DS metric that captures the deviation of irradiated cell populations from non-irradiated controls in a multidimensional biomarker space, thus showing a duality behaviour with respect to the gold-standard dose-response profile. In fact, the DS curve consistently displayed the hallmark properties of radiobiological dose–response relationships, i.e. monotonicity, nonlinearity, and a defined baseline at zero dose. Importantly, when normalized for assessment purposes, the DS curve showed strong correlations with gold-standard clonogenic survival curves across three model cancer cell lines (neuroblastoma SHSY5Y cells, neuroblastoma SKNBE2 cells, and breast cancer MCF7 cells), with Pearson correlation coefficients ranging from 0.94 to 0.99. These results confirm that HTFC-derived DS curves can serve as reliable label-free alternative to standard clonogenic assays.

Compared to conventional assays, the HTFC approach provides several critical advantages. First, its label-free aspect eliminates the need for exogenous dyes, thus avoiding issues such as photobleaching, phototoxicity, or interference with downstream analyses. Second, it is rapid as it enables acquisition and analysis of hundreds/thousands of single cells within 24 hours after irradiation, whereas clonogenic assays typically require 1-2 weeks. Third, it is operator-independent, providing an objective, automated readout that reduces bias inherent in colony counting. Above all, by extracting a rich set of intracellular biophysical markers, HTFC goes beyond survival metrics, offering detailed insights into 3D morphological and RI changes associated with radiation damage at subcellular level.

Unlike conventional biosensors that rely on indirect readouts based on molecular probes or fluorescent labels, HTFC combines physical optics with flow cytometry to directly measure cell states in a minimally invasive manner. In this study, we observed a strong correlation between the DS curve and the standard SF curve (>90%). In classical radiobiology, however, cell survival following irradiation is typically modelled by fitting SF data with the linear–quadratic (LQ) model [48] or other advanced approaches [49]. For example, the LQ formalism expresses the logarithm of the SF as the sum of a linear term (αD), representing irreparable lethal damage proportional to dose, and a quadratic term (βD^2^), which accounts for sublethal damage that becomes lethal when accumulated or misrepaired. The α/β ratio is widely regarded as a key indicator of radiosensitivity to guide clinical dose-fractionation strategies, distinguishing tissues or tumours with predominantly linear (early responding) vs. quadratic (late responding) damage responses [50]. Parameters of the LQ model are shown as example in Figs. S3(a-c), corresponding to the SF curves of the three model cell lines analyzed in this study. However, the same radiobiological models cannot be directly applied to the DS dose-response curve, since its biological meaning differs from that of the SF curve. Clonogenic assays quantify the long-term reproductive survival of irradiated cells, measuring the fraction of cells that retain the ability to proliferate after radiation exposure. In contrast, HTFC provides an early assessment, as it quantifies radiation-induced alterations in the biophysical properties of cells that remain viable 24 hours after treatment. Thus, while SF reflects the ultimate proliferative capacity of a population, DS captures short-term phenotypic and structural changes in surviving cells. Therefore, future work will focus on identifying and calibrating the most suitable model curves to translate DS-derived biological responses into precise dose-response parameters across different cell lines and tissue types. Additionally, ongoing studies will aim to extend and validate this label-free approach using cells derived from patient biopsies.

In conclusion, we have demonstrated that HTFC-derived DS curves offer a label-free, rapid, and operator-independent assay that could in future improve the classical and well assessed clonogenic survival analyses. This study represents a significant advance, opening the way to predictive biosensors for personalized RT in the future. In fact, these preliminary results indicate that HTFC has the potential to serve as a next-generation biosensor, capable of rapidly profiling tumour radiosensitivity from patient-derived samples and supporting the development of precision oncology strategies in the field of personalized RT.

## Methods

### Sample preparation

Neuroblastoma SHSY5Y and SKNBE2 cells, as well as breast cancer MCF7 cells, were cultured in Falcon^®^ T25 flasks under standard conditions. SHSY5Y cells were maintained in Dulbecco’s Modified Eagle Medium (DMEM, Avantor, VWR) supplemented with 20% fetal bovine serum (FBS, Avantor, VWR), 1% penicillin-streptomycin (P/S, biowest), and 1% L-glutamine (Sigma, St. Louis, MO, USA). SKNBE2 cells were cultured in a 1:1 mixture of DMEM and Ham’s F-12 nutrient mixture (Ham’s F12, Avantor, VWR) supplemented with 10% FBS, 1% P/S, and 1% L-glutamine. MCF7 cells were maintained in DMEM supplemented with 10% FBS, 1% P/S.

All the samples for all the cell lines were prepared at a concentration of ≥ 300,000 cells/ml to meet HTFC experimental requirements. Cell vitality and proliferation were monitored daily using the LUNA-II™ Automated Brightfield Cell Counter. Cultures were incubated at 37 °C in a humidified atmosphere containing 5% CO2. The cell concentrations for the clonogenic assay varied on the doses.

### Irradiation experiments

Neuroblastoma SHSY5Y cells, neuroblastoma SKNBE2 cells, and breast cancer MCF7 cells were exposed to X-rays administered through conventional external beam radiotherapy at the IRCCS Istituto Nazionale Tumori Fondazione Pascale in Naples (Italy). The irradiation of samples was conducted using the Versa HD clinical LINAC system by ELEKTA, which utilizes a 6 MV electron beam to generate X-rays. Different radiation doses were delivered to the cell cultures using 3D conformal radiotherapy (3D-CRT). In particular, 0.5 Gy, 1 Gy, 2 Gy, and 4 Gy were used with SHSY5Y cells, while 0.25 Gy, 0.5 Gy, 1 Gy, 1.5 Gy, and 2 Gy were used with SKNBE2 and MCF7 cells. The 3D-CRT treatment plans were developed with Elekta’s Monaco v5.11.03 treatment planning system (TPS), which combines Monte Carlo dose calculation accuracy with robust optimization tools to deliver high-quality radiotherapy treatment. Falcon^®^ T25 cell culture flasks were placed in a uniform 20×20 cm^2^ square field and positioned between two plexiglass plates (ScandiDos Delta-4 Calibration Phantom): the first plate (3 cm thick) was used to simulate the dose buildup effect for shallow penetration depths, while the second plate (5 cm thick) was included to stop backscattered radiation. The cell samples were irradiated from opposing fields via a 180° gantry rotation, delivering X-rays at a fixed dose rate of 200 monitor units (MU) per minute. This experimental setup allowed for the uniform administration of the prescribed doses at the cellular level. Following clinical radiotherapy exposure, the cells were stored in an incubator at 37°C in a 5% CO2 atmosphere, in preparation for subsequent HTFC analyses (24 hours post-irradiation) and clonogenic survival assays (1-2 weeks post-irradiation).

### Clonogenic assay

The clonogenic assay, the gold standard for measuring cellular radiosensitivity, was used to assess radiation-induced cell death [47]. The assay defines a cell as surviving irradiation if it retains its proliferative potential and forms a colony of at least 50 cells. Cells were seeded at different densities the day before the irradiation, with the concentration optimized according to the radiation dose. Following exposure, the flasks were returned to the incubator. After 1-2 weeks of incubation (9 days for neuroblastoma SKNBE2 cells, 14 days for breast cancer MCF7 cells, and 14 days for neuroblastoma SHSY5Y cells), cells were fixed and the resulting colonies were quantified. The surviving fraction SF for a given dose D was calculated as

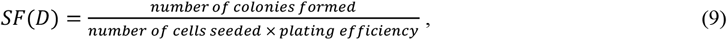

where the plating efficiency PE was calculated as

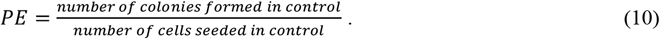

The SF data were finally fitted using an LQ model, i.e. [48]

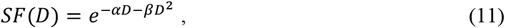

as also shown in Fig. S3.

For each dose, three flasks were seeded at the same density and irradiated to allow replication of the SF calculation.

### HTFC experiments

Hologram acquisition was carried out using a Mach–Zehnder interferometer with symmetric optical paths [35], as illustrated in Fig. 6(a). A collimated light source (SuperK FIANIUM, 495 nm, FWHM ≈ 4 nm) was split by beam splitter BS1 into an object and a reference beam. The object beam interacts with cells flowing through a microfluidic channel (PMMA, 1000×200 µm^2^), and is collected by a microscope objective (MO1, Zeiss 63×/0.95 numerical aperture), followed by a tube lens L1 (150 mm focal length) and a telescope system composed of lenses L2 (150 mm focal length) and L5 (200 mm focal length). The reference beam travels along a separate optical path containing equivalent components: a microscope objective (MO2, Zeiss 63X/1.3 numerical aperture, dry) and tube lens L3 (150 mm focal length), followed by a second telescope with lenses L4 (150 mm focal length) and L5. The two beams are recombined at a second beam splitter (BS2). Temporal coherence is maintained using delay lines implemented with mirrors M1, M2 and M6, M7 in the object and reference arms, respectively. Spatial coherence is ensured by introducing a diffraction grating (Newport Moiré Grating, 80 lp/mm) in the reference arm and an iris diaphragm (I) to select only the first diffraction order. This choice assures homogeneity of the interference fringes across the whole field of view. The resulting off-axis interference pattern is captured by a CMOS camera (Genie Nano-CXP, 5120×5120, 4.5 µm pixel size).

**Figure 6.**
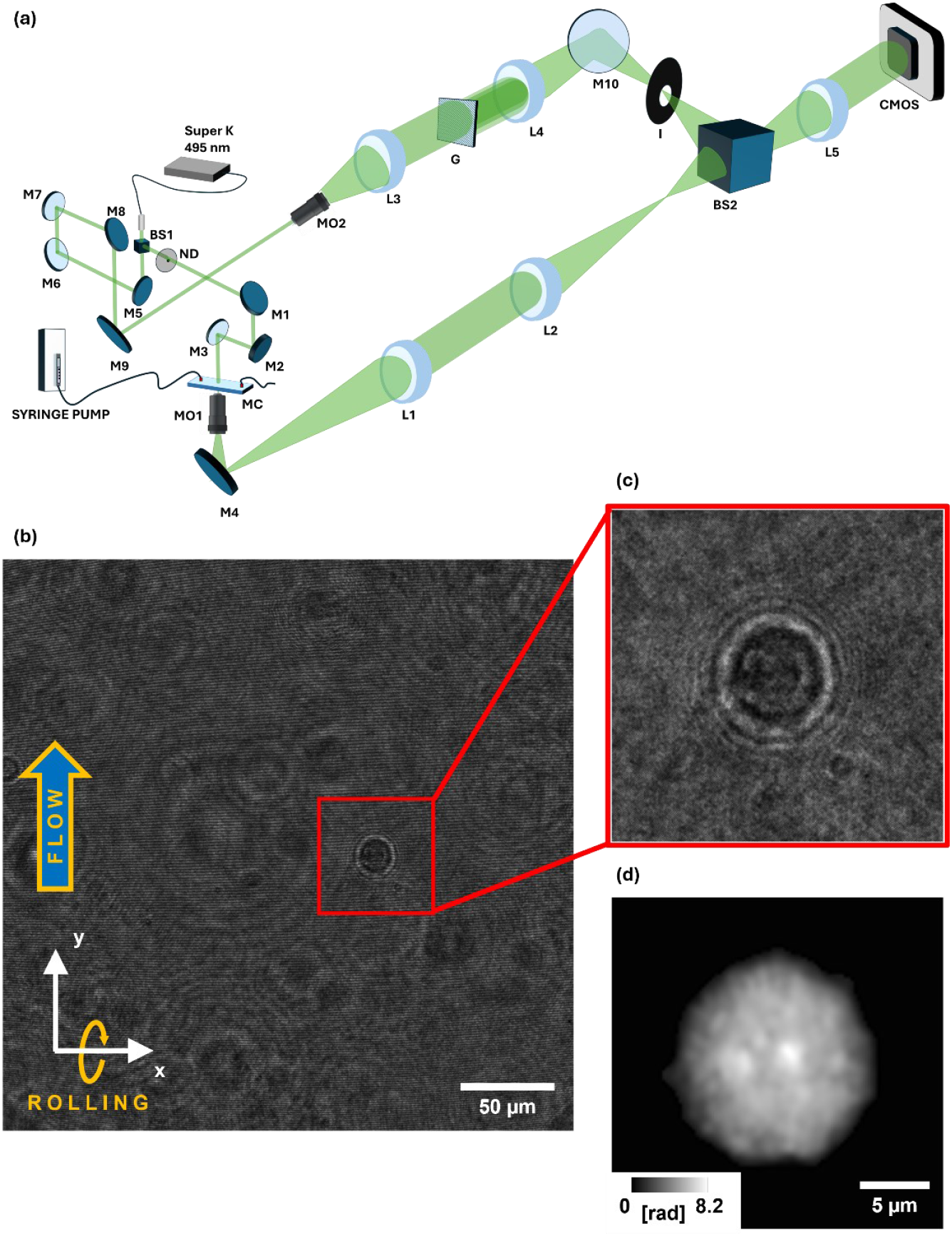
HTFC experimental system and numerical reconstruction. **(a)** Sketch of the opto-fluidic recording system. SuperK, laser-light source; BS, beam splitters; ND, neutral density filter; M, mirrors; MO, microscope objectives; L, lenses; G, diffracti on grating; I, iris diaphragm. **(b)** Hologram taken from a recorded video sequence of control neuroblastoma SKNBE2 cells. **(c)** Sub-hologram highlighted in (b). **(d)** QPM reconstructed from the sub-hologram in (c).

To acquire multiple views of individual cells, hydrodynamic rotation was induced by positioning cells off-center in the microfluidic channel while maintaining a controlled flow rate of 75 nL/s using a CETONI neMESYS 290N syringe pump [29-35]. According to the reference system in Fig. 6(b), cells flow along the y-axis and rotate around the x-axis. From each hologram of the recorded video sequence (5120×5120 corresponding to 305×305 µm^2^), cells were tracked to extract squared sub-holograms containing each rotating cell (1024×1024 corresponding to 61×61 µm^2^), as shown in Fig. 6(b). For each sub-hologram (see Fig. 6(c)), the real diffraction order was isolated via band-pass Fourier filtering exploiting the off-axis configuration [17]. The Angular Spectrum propagation method [17] was used to determine the optimal focus distance for each cell, minimizing the Tamura coefficient [51]. Phase-contrast images were obtained from the argument of the in-focus complex field, followed by phase aberration correction through subtraction of a reference hologram [52] and phase denoising based on the windowed Fourier transform filter [53]. Finally, phase unwrapping based on the PUMA algorithm [54] was applied to generate QPMs for each sub-hologram (201×201 corresponding to 24×24 µm^2^), as displayed in Fig. 6(d). For each cell, the stack of QPMs containing its rotation was employed to estimate the unknown rolling angles [55], and, finally, its 3D RI tomogram was reconstructed using the Filtered Back Projection algorithm [29-33,56], as reported in Fig. 1(a).

## Data availability

The data that support the findings of this study are available from the corresponding authors upon request.

## Acknowledgments

This work was supported by project PRIN 2022 PNRR - Flow-cytometry ImaGing by Holographic tomography for predicting TUMor control in Oncology patients treated with Radiotheraphy (FIGHT-TUMOR), Prot. P2022ATE2J – funded by the Italian Ministry of University & Research in the framework of Next Generation EU.

## Additional information

Supplementary data to this article can be found in the Supplementary Information.

