## Supplementary Information for "Establishing label-free quantitative X-ray dose-response profiling by Holo-Tomographic Flow Cytometry"

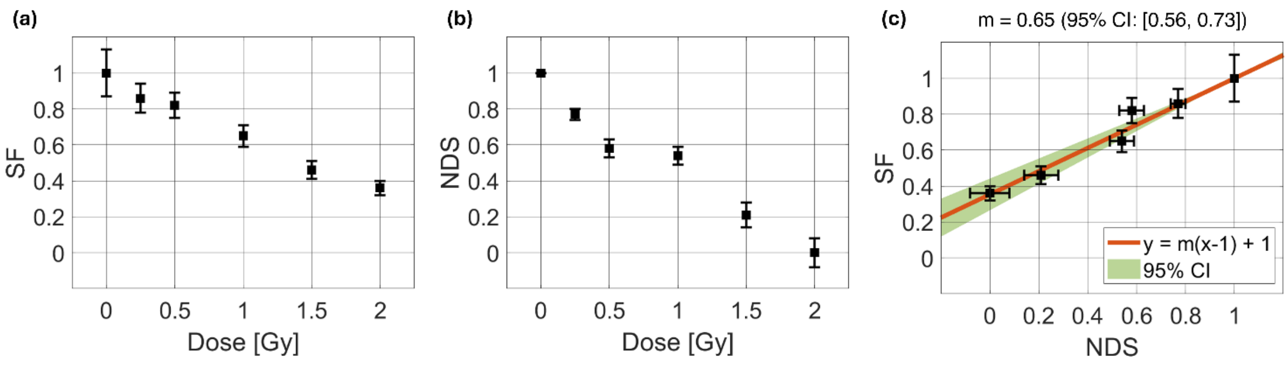


**Figure S1. Assessment of the label-free X-ray dose-response curve for the breast cancer MCF7 cell line. (a,b)** SF and NDS curves, respectively**. (c)** Correlation between SF curve in (a) and NDS curve in (b).

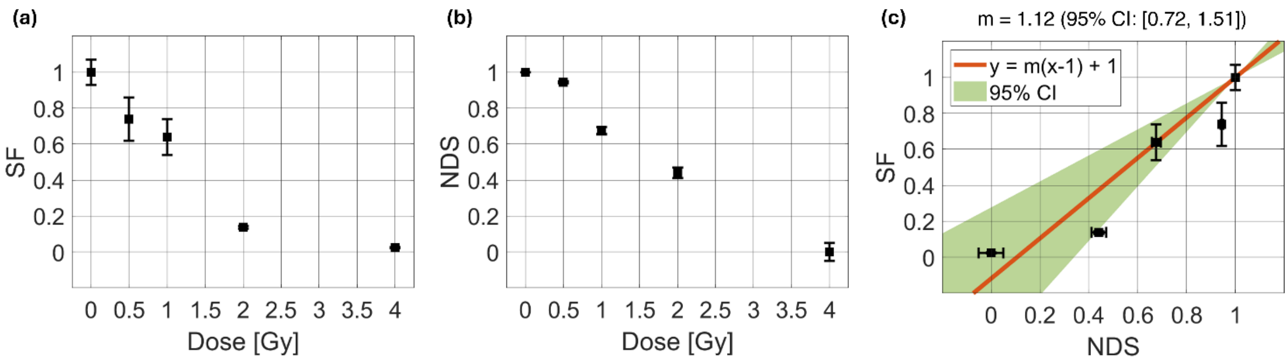


**Figure S2. Assessment of the label-free X-ray dose-response curve for the neuroblastoma SHSY5Y cell line. (a,b)** SF and NDS curves, respectively**. (c)** Correlation between SF curve in (a) and NDS curve in (b).

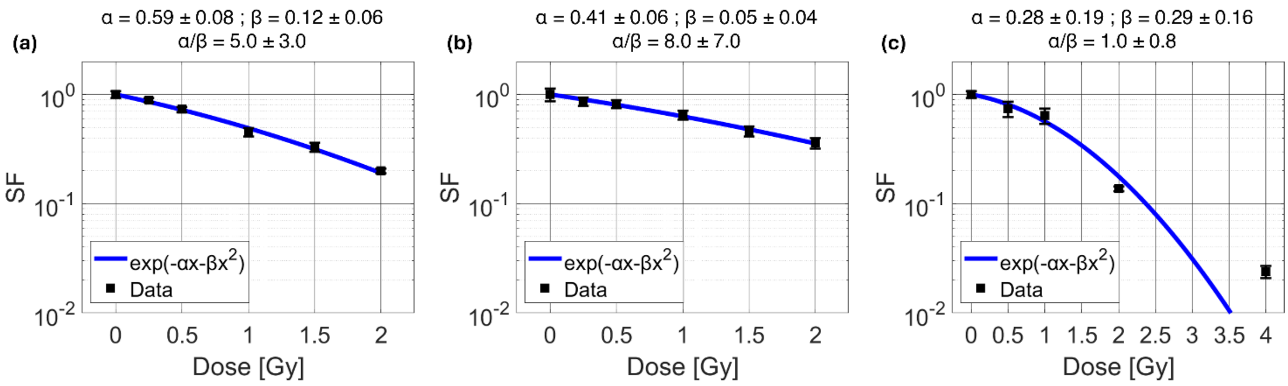


**Figure S3. Fitting of the SF curves (black dots) through the LQ model (blue curve), reported in semilogarithmic scale. (a)** Neuroblastoma SKNBE2 cell line. **(b)** Breast cancer MCF7 cell line. **(c)** Neuroblastoma SHSY5Y cell line. Parameters of the LQ model are reported at the top.

| Dose [Gy] | # Tomograms |
|---|---|
| 0 Gy | 109 |
| 0.25 Gy | 127 |
| 0.5 Gy | 83 |
| 1 Gy | 87 |
| 1.5 Gy | 87 |
| 2 Gy | 97 |
| | 590 |

**Table S1.** Dataset of 3D RI tomograms about neuroblastoma SKNBE2 cells collected by the HTFC system.

| Dose | DS (or FDR) | NDS | SF |
|---|---|---|---|
| 0 Gy | 0 | 1 | 1.00 ± 0.07 |
| 0.25 Gy | 0.0046 ± 0.0013 | 0.85 ± 0.04 | 0.89 ± 0.04 |
| 0.5 Gy | 0.014 ± 0.002 | 0.55 ± 0.08 | 0.73 ± 0.04 |
| 1 Gy | 0.025 ± 0.003 | 0.19 ± 0.12 | 0.45 ± 0.04 |
| 1.5 Gy | 0.029 ± 0.003 | 0.06 ± 0.13 | 0.33 ± 0.03 |
| 2 Gy | 0.031 ± 0.003 | 0.00 ± 0.14 | 0.20 ± 0.01 |

**Table S2.** DS (or FDR), NDS, and SF values of the neuroblastoma SKNBE2 cell line.

| Dose [Gy] | # Tomograms |
|---|---|
| 0 Gy | 44 |
| 0.25 Gy | 89 |
| 0.5 Gy | 116 |
| 1 Gy | 115 |
| 1.5 Gy | 75 |
| 2 Gy | 89 |
| | 528 |

**Table S3.** Dataset of 3D RI tomograms about breast cancer MCF7 cells collected by the HTFC system.

| Dose | DS (or FDR) | NDS | SF |
|---|---|---|---|
| 0 Gy | 0 | 1 | 1.00 ± 0.13 |
| 0.25 Gy | 0.047 ± 0.006 | 0.77 ± 0.03 | 0.86 ± 0.08 |
| 0.5 Gy | 0.085 ± 0.008 | 0.58 ± 0.05 | 0.82 ± 0.07 |
| 1 Gy | 0.094 ± 0.008 | 0.54 ± 0.05 | 0.65 ± 0.06 |
| 1.5 Gy | 0.160 ± 0.011 | 0.21 ± 0.07 | 0.46 ± 0.05 |
| 2 Gy | 0.203 ± 0.012 | 0.00 ± 0.08 | 0.36 ± 0.04 |

**Table S4.** DS (or FDR), NDS, and SF values of the breast cancer MCF7 cell line.

| Dose [Gy] | # Tomograms |
|---|---|
| 0 | 38 |
| 0.5 | 133 |
| 1 | 50 |
| 2 | 38 |
| 4 | 24 |
| | 283 |

**Table S5.** Dataset of 3D RI tomograms about neuroblastoma SHSY5Y cells collected by the HTFC system.

| Dose | DS (or FDR) | NDS | SF |
|---|---|---|---|
| 0 Gy | 0 | 1 | 1.00 ± 0.07 |
| 0.5 Gy | 0.080 ± 0.007 | 0.944 ± 0.005 | 0.74 ± 0.12 |
| 1 Gy | 0.46 ± 0.02 | 0.676 ± 0.018 | 0.64 ± 0.10 |
| 2 Gy | 0.79 ± 0.03 | 0.44 ± 0.03 | 0.137 ± 0.007 |
| 4 Gy | 1.42 ± 0.05 | 0.00 ± 0.05 | 0.024 ± 0.003 |

**Table S6.** DS (or FDR), NDS, and SF values of the neuroblastoma SHSY5Y cell line.

| | Biomarker | Description |
|---|---|---|
| 1 | RI Mean - Cell | Average RI value of the cell |
| 2 | RI Standard Deviation - Cell | Standard deviation of the RI values of the cell |
| 3 | RI Coefficient of Variation - Cell | Ratio between the RI Standard Deviation and the RI Mean of the cell |
| 4 | RI Maximum - Cell | Maximum RI value of the cell |
| 5 | RI Interquartile Range - Cell | Difference between the III and the I quartile of the RI distribution of the cell |
| 6 | Volume - Cell | Geometric volume of the cell |
| 7 | Dry Mass - Cell | Mass of the cell without considering its aqueous content |
| 8 | Sphericity - Cell | Ratio of the surface area of a sphere with the same volume to the cell's surface area |
| 9 | Solidity - Cell | Ratio between the volume of the cell and the volume of its convex hull |
| 10 | Extent - Cell | Ratio between the volume of the cell and the volume of its bounding box |
| 11 | RI Mean - Nucleus | Average RI value of the nucleus |
| 12 | RI Standard Deviation - Nucleus | Standard deviation of the RI values of the nucleus |
| 13 | RI Coefficient of Variation - Nucleus | Ratio between the RI Standard Deviation and the RI Mean of the nucleus |
| 14 | RI Maximum - Nucleus | Maximum RI value of the nucleus |
| 15 | RI Interquartile Range - Nucleus | Difference between the III and the I quartile of the RI distribution of the nucleus |
| 16 | Volume - Nucleus | Geometric volume of the nucleus |
| 17 | Dry Mass - Nucleus | Mass of the nucleus without considering its aqueous content |
| 18 | Sphericity - Nucleus | Ratio of the surface area of a sphere with the same volume to the nucleus surface area |
| 19 | Solidity - Nucleus | Ratio between the volume of the cell and the volume of its convex hull |
| 20 | Extent - Nucleus | Ratio between the volume of the nucleus and the volume of its bounding box |
| 21 | RI Mean - Cytoplasm | Average RI value of the cytoplasm |
| 22 | RI Standard Deviation - Cytoplasm | Standard deviation of the RI values of the cytoplasm |
| 23 | RI Coefficient of Variation - Cytoplasm | Ratio between the RI Standard Deviation and the RI Mean of the cytoplasm |
| 24 | RI Maximum - Cytoplasm | Maximum RI value of the cytoplasm |
| 25 | RI Interquartile Range - Cytoplasm | Difference between the III and the I quartile of the RI distribution of the cytoplasm |
| 26 | Volume - Cytoplasm | Geometric volume of the cytoplasm |
| 27 | Dry Mass - Cytoplasm | Mass of the cytoplasm without considering its aqueous content |
| 28 | RI Mean – Nucleus/Cytoplasm | Ratio between the RI Mean of the nucleus and the cytoplasm |
| 29 | RI Standard Deviation - Nucleus/Cytoplasm | Ratio between the RI Standard Deviation of the nucleus and the cytoplasm |
| 30 | RI Coefficient of Variation - Nucleus/Cytoplasm | Ratio between the RI Coefficient of Variation of the nucleus and the cytoplasm |
| 31 | RI Maximum - Nucleus/Cytoplasm | Ratio between the RI Maximum of the nucleus and the cytoplasm |
| 32 | RI Interquartile Range - Nucleus/Cytoplasm | Ratio between the RI Interquartile Range of the nucleus and the cytoplasm |
| 33 | Volume - Nucleus/Cytoplasm | Ratio between the Volume of the nucleus and the cytoplasm |
| 35 | Dry Mass - Nucleus/Cytoplasm | Ratio between the Dry Mass of the nucleus and the cytoplasm |
| 35 | Sphericity – Nucleus/Cell | Ratio between the Sphericity of the nucleus and the cell |
| 36 | Solidity - Nucleus/Cell | Ratio between the Solidity of the nucleus and the cell |
| 37 | Extent - Nucleus/Cell | Ratio between the Extent of the nucleus and the cell |
| 38 | Nucleus-Cell Normalized Distance | Euclidean distance between the nucleus centroid and the cell centroid normalized to the cell equivalent radius, that is defined as the radius of a sphere having the same volume as the cell |
| 39 | Nucleus-Cell Normalized Solid Angle | Solid angle subtending the nucleus from the cell centroid, normalized to 4π |

**Table S7.** List and description of the 3D biophysical markers used to characterize the HTFC dataset.
